## Supplementary tables and figures for "Multiple Functions of Cerebello-Thalamic Neurons in Learning and Offline Consolidation of a Motor Skill in mice"

| Site | Group 1 | Group 2 | n1 | n2 | Test | Alternative | Statistic | p-value | Adj. p-value | Sig | Effect Size |
|---|---|---|---|---|---|---|---|---|---|---|---|
| DCN | DREADD+CNO | DREADD+Saline | 75 | 77 | Wilcoxon | two.sided | 2119 | 0.005 | 0.019 | * | 0.23 |
| | DREADD+CNO | Sham+CNO | 75 | 69 | | | 1518 | 0.0000191 | 0.000115 | *** | 0.356 |
| | DREADD+CNO | Sham+Saline | 75 | 39 | | | 899 | 0.000772 | 0.004 | ** | 0.315 |
| | DREADD+Saline | Sham+CNO | 77 | 69 | | | 2429 | 0.374 | 1 | | 0.0738 |
| | DREADD+Saline | Sham+Saline | 77 | 39 | | | 1438 | 0.713 | 1 | | 0.0345 |
| | Sham+CNO | Sham+Saline | 69 | 39 | | | 1493 | 0.347 | 1 | | 0.0908 |

Supplementary Table 1: Statistics for Fig 1 g.

| Site | Group | Factor | ANOVA | p-value | Sig. | Test | Group 1 | Group 2 | Statistic | CI 2.5% | CI 97.5% | p-value | Cohen's D | Sig. |
|---|---|---|---|---|---|---|---|---|---|---|---|---|---|---|
| DCN | DREADD | Treatment | F(1,106)=7.96 | 0.006 | ** | | | | | | | | | |
| | | Speed | F(1,106)=201.619 | <0.001 | *** | | | | | | | | | |
| | | Treatment:Speed | F(1,106)=1.039 | 0.31 | n.s. | post-hoc | 5 rpm-SAL | 5 rpm-CNO | -0.449 | -12.7 | 8.25 | 0.659 | -0.181 | n.s. |
| | | | | | | t-test | 10 rpm-SAL | 10 rpm-CNO | -1.37 | -35.5 | 7.91 | 0.193 | -0.542 | n.s. |
| | | | | | | | 15 rpm-SAL | 15 rpm-CNO | -1.41 | -84.2 | 16.3 | 0.174 | -0.582 | n.s. |
| | | | | | | | 20 pm-SAL | 20 rpm-CNO | -1.76 | -106 | 9.76 | 0.097 | -0.775 | n.s. |
| | | | | | | | 25 rpm-SAL | 25 rpm-CNO | -1.78 | -30.6 | 3.46 | 0.106 | -0.828 | n.s. |
| | Sham | Treatment | F(1,76)=1.836 | 0.179 | n.s. | | | | | | | | | |
| | | Speed | F(1,76)=152.296 | <0.001 | *** | | | | | | | | | |
| | | Treatment:Speed | F(1,76)=0.109 | 0.742 | n.s. | | | | | | | | | |

Supplementary Table 2: Statistics for Fig 1 h.

| Site | Time of treatment | Day | Factor | ANOVA | p-value | Sig. |
|---|---|---|---|---|---|---|
| DCN | During task | 1 | Treatment | F(3,45)=0.9452 | 0.426 | n.s. |
| | | 2 | | F(3,45)=4.021 | 0.013 | * |
| | | 3 | | F(3,45)=4.212 | 0.01 | * |
| | | 4 | | F(3,45)=6.601 | <0.001 | *** |
| | | 5 | | F(3,45)=7.899 | <0.001 | *** |
| | | 6 | | F(3,45)=13.33 | <0.001 | *** |
| | | 7 | | F(3,45)=7.741 | <0.001 | *** |
| | After task | 1 | | F(3,30)=0.414 | 0.744 | n.s. |
| | | 2 | | F(3,30)=6.215 | 0.002 | ** |
| | | 3 | | F(3,30)=3.868 | 0.019 | * |
| | | 4 | | F(3,30)=2.26 | 0.101 | n.s. |
| | | 5 | | F(3,30)=1.413 | 0.258 | n.s. |
| | | 6 | | F(3,30)=0.256 | 0.856 | n.s. |
| | | 7 | | F(3,30)=1.971 | 0.1396 | n.s. |

Supplementary Table 3: Statistics for Fig 1 ij part1.

| Site | Time of treatment | Comparaison | Moment 1 | Moment 2 | Test | DREADD + CNO | DREADD + SAL | Sham + CNO | Sham + SAL |
|---|---|---|---|---|---|---|---|---|---|
| DCN | During task | overnight | Trial 7 Day 1 | Trial 1 Day2 | posthoc on | 0.031 * | 1 | 1 | 0.9 |
| | | | Trial 7 Day 2 | Trial 1 Day3 | estimated | 0.223 | 0.58 | 0.15 | 0.94 |
| | | | Trial 7 Day 3 | Trial 1 Day4 | means | 0.037 * | 1 | 0.29 | 1 |
| | | | Trial 7 Day 4 | Trial 1 Day5 | | 0.071 | 1 | 0.39 | 0.27 |
| | | | Trial 7 Day 5 | Trial 1 Day6 | | 0.223 | 0.26 | 1 | 1 |
| | | | Trial 7 Day 6 | Trial 1 Day7 | | 0.223 | 1 | 1 | 1 |
| | After task | overnight | Trial 7 Day 1 | Trial 1 Day2 | | <0.001 *** | 1 | 0.933 | 1 |
| | | | Trial 7 Day 2 | Trial 1 Day3 | | <0.001 *** | 1 | 0.043 * | 1 |
| | | | Trial 7 Day 3 | Trial 1 Day4 | | 0.305 | 1 | 0.596 | 1 |
| | | | Trial 7 Day 4 | Trial 1 Day5 | | -0.076 | 1 | 0.362 | 1 |
| | | | Trial 7 Day 5 | Trial 1 Day6 | | 0.463 | 1 | 0.247 | 1 |
| | | | Trial 7 Day 6 | Trial 1 Day7 | | 0.405 | 0.26 | 0.633 | 1 |

Supplementary Table 4: Statistics for Fig ij part2.

| Site | Estimate | Comparison | n | Estimate | Test | Alternative | Statistic | p-value | Sig. | CI 2.5% | CI 97.5% | Effect Size |
|---|---|---|---|---|---|---|---|---|---|---|---|---|
| DCN pooled Ctrl. | Consolidated learning | day 2 - day 1 | 61 | 45.8 | Wilcoxon | two.sided | 1833 | 0.000000000188 | *** | 35.3 | 55.6 | 0.816 |
| | | day 3 - day 2 | 61 | 22.2 | | | 1603 | 0.00000237 | *** | 14.7 | 29 | 0.605 |
| | | day 4 - day 3 | 61 | 13.4 | | | 1219 | 0.0499 | * | 0.0714 | 26.1 | 0.252 |
| | | day 5 - day 4 | 61 | 7.07 | | | 1148 | 0.147 | n.s. | -2.77 | 17.7 | 0.186 |
| | | day 6 - day 5 | 61 | 4.14 | | | 1011 | 0.482 | n.s. | -7.29 | 16.1 | 0.091 |
| | | day 7 - day 6 | 61 | 4.5 | | | 1063 | 0.401 | n.s. | -6.84 | 14.8 | 0.108 |

Supplementary Table 5: Statistics for Fig 1 l.

| Site | Experiment | Value | ANOVA Group | p-value | Sig. |
|---|---|---|---|---|---|
| DCN | Footprint | Linear movement | F(3,31)=0.887 | 0.459 | n.s. |
| | | Sigma | F(3,31)=0.103 | 0.958 | n.s. |
| | | Alternate coefficient | F(3,31)=0.476 | 0.701 | n.s. |
| | Grid test | Latency | F(3,33)=0.305 | 0.822 | n.s. |
| | Horizontal Bar | Latency | F(3,33)=0.826 | 0.489 | n.s. |
| | Vertical Pole | Latency | F(3,33)=0.968 | 0.42 | n.s. |

Supplementary Table 6: Statistics for Sup Fig 2 ab.

| Site | Group | Day | Factor | ANOVA | p-value | Sig. | Group 1 | Group 2 | Statistic | CI 2.5% | CI 97.5% | p-value | Cohen's D | Sig. |
|---|---|---|---|---|---|---|---|---|---|---|---|---|---|---|
| DCN | DREADD | day 1 | Treatment | F(1,30)=2.102 | 0.157 | n.s. | | | | | | | | |
| | | | Treatment:Moment | F(1,30)=0.87 | 0.358 | n.s. | | | | | | | | |
| | | day 4 | Treatment | F(1,30)=0.493 | 0.488 | n.s. | | | | | | | | |
| | | | Treatment:Moment | F(1,30)=0.595 | 0.446 | n.s. | | | posthoc | | | | | |
| | | day 7 | Treatment | F(1,30)=0.131 | 0.72 | n.s. | | | ttest | | | | | |
| | | | Treatment:Moment | F(1,30)=8.61 | 0.006 | ** | OF1 (before SAL) | OF1 (before CNO) | 1.818 | -0.103 | 1.1 | 0.095 | 0.848 | n.s. |
| | | | | | | | OF2 (after SAL) | OF2 (after CNO) | -1.384 | -1.63 | 0.351 | 0.188 | -0.657 | n.s. |
| | Sham | day 1 | Treatment | F(1,32)=5.663 | 0.023 | * | | | | | | | | |
| | | | Treatment:Moment | F(1,32)=0.078 | 0.782 | n.s. | OF1 (before SAL) | OF1 (before CNO) | -0.656 | -1.91 | 1.01 | 0.521 | -0.309 | n.s. |
| | | | | | | | OF2 (after SAL) | OF2 (after CNO) | -0.543 | -1.76 | 1.04 | 0.595 | -0.256 | n.s. |
| | | day 4 | Treatment | F(1,32)=3.874 | 0.058 | n.s. | | | | | | | | |
| | | | Treatment:Moment | F(1,32)=0.153 | 0.698 | n.s. | | | | | | | | |
| | | day 7 | Treatment | F(1,32)=0.761 | 0.39 | n.s. | | | | | | | | |
| | | | Treatment:Moment | F(1,32)=1.4 | 0.245 | n.s. | | | | | | | | |

Supplementary Table 7: Statistics for Sup Fig 2 c.

| Comparison | Group | Intcp Ctrl | Slope Ctrl | Median dist. Ctrl-Deming | n Ctrl | Median dist. Grp-Deming | n Group | Est. diff. | CI 2.5% | CI 97.5% | Statistics Wilcoxon | p-value | Sig. | Eff. Size |
|---|---|---|---|---|---|---|---|---|---|---|---|---|---|---|
| Learning | DREADD+CNO during task | 109 | -0.801 | -1.77 | 244 | -20.6 | 56 | -14.8 | -23.2 | -6.35 | 4821 | 0.000594 | *** | 0.198 |
| Consolidation | DREADD+CNO during task | -9.86 | 1.04 | -1.33 | 244 | -13.0 | 56 | -12.1 | -19.9 | -4.76 | 4973 | 0.0015 | ** | 0.183 |
| Learning | DREADD+CNO after task | 109 | -0.801 | -1.77 | 244 | -15.2 | 32 | -10 | -20 | 0.247 | 3093 | 0.0562 | n.s. | 0.115 |
| Consolidation | DREADD+CNO after task | -9.86 | 1.04 | -1.33 | 244 | -17.4 | 32 | -14.9 | -24.1 | -6.21 | 2512 | 0.00105 | ** | 0.197 |

Supplementary Table 8: Statistics for Fig 2de. Deming regression.

| Group | Time of treatment | Day | ANOVA Treatnent | p-value | Sig. | ANOVA Treatnent:Trial | p-value | Sig. |
|---|---|---|---|---|---|---|---|---|
| CN – CL DREADD | During task | 1 | F(1,18)=1.61 | 0.221 | n.s. | F(6,108)=0.5894 | 0.7382 | n.s. |
| | | 2 | F(1,18)=7.548 | 0.013 | * | F(6,108)=0.4718 | 0.828 | n.s. |
| | | 3 | F(1,18)=5.057 | 0.037 | * | F(6,108)=1.023 | 0.414 | n.s. |
| | | 4 | F(1,18)=10.28 | 0.004 | ** | F(6,108)=0.2491 | 0.9587 | n.s. |
| | | 5 | F(1,18)=8.577 | 0.008 | ** | F(6,108)=0.9642 | 0.453 | n.s. |
| | | 6 | F(1,18)=15.97 | <0.001 | *** | F(6,108)=0.5614 | 0.7602 | n.s. |
| | | 7 | F(1,18)=8.386 | 0.009 | ** | F(6,108)=0.5056 | 0.8029 | n.s. |
| | After task | 1 | F(1,17)=0.3552 | 0.559 | n.s. | F(6,102)=2.68 | 0.0186 | # |
| | | 2 | F(1,17)=0.5855 | 0.454 | n.s. | F(6,102)=1.179 | 0.3235 | n.s. |
| | | 3 | F(1,17)=4.418 | 0.051 | n.s. | F(6,102)=3.069 | 0.0084 | ## |
| | | 4 | F(1,17)=1.494 | 0.238 | n.s. | F(6,102)=0.773 | 0.5929 | n.s. |
| | | 5 | F(1,17)=1.3 | 0.27 | n.s. | F(6,102)=4.524 | <0.001 | ### |
| | | 6 | F(1,17)=0.01609 | 0.901 | n.s. | F(6,102)=0.3709 | 0.8959 | n.s. |
| | | 7 | F(1,17)=1.585 | 0.225 | n.s. | F(6,102)=1.124 | 0.3539 | n.s. |
| CN – VAL DREADD | During task | 1 | F(1,19)=1.914 | 0.1826 | n.s. | F(6,114)=0.9818 | 0.441 | n.s. |
| | | 2 | F(1,19)=2.368 | 0.1403 | n.s. | F(6,114)=1.507 | 0.1819 | n.s. |
| | | 3 | F(1,19)=0.3523 | 0.5598 | n.s. | F(6,114)=0.7044 | 0.6467 | n.s. |
| | | 4 | F(1,19)=4.968 | 0.03809 | * | F(6,114)=0.7656 | 0.5985 | n.s. |
| | | 5 | F(1,19)=5.973 | 0.0245 | * | F(6,114)=0.5808 | 0.745 | n.s. |
| | | 6 | F(1,19)=17.28 | 0.000536 | *** | F(6,114)=1.797 | 0.1058 | n.s. |
| | | 7 | F(1,19)=9.661 | 0.006 | ** | F(6,114)=1.223 | 0.2996 | n.s. |
| | After task | 1 | F(1,16)=3.824 | 0.06821 | n.s. | F(6,96)=1.381 | 0.23 | n.s. |
| | | 2 | F(1,16)=2.827 | 0.1121 | n.s. | F(6,96)=2.635 | 0.02078 | # |
| | | 3 | F(1,16)=0.03395 | 0.8561 | n.s. | F(6,96)=2.015 | 0.07095 | n.s. |
| | | 4 | F(1,16)=0.06177 | 0.8069 | n.s. | F(6,96)=3.911 | 0.001543 | ## |
| | | 5 | F(1,16)=0.006183 | 0.938 | n.s. | F(6,96)=2.844 | 0.01361 | # |
| | | 6 | F(1,16)=0.1843 | 0.6734 | n.s. | F(6,96)=4.661 | <0.001 | ### |
| | | 7 | F(1,16)=2.812 | 0.113 | n.s. | F(6,96)=1.507 | 0.1839 | n.s. |

Supplementary Table 9: Statistics for Fig 3 bcef.

| Comparison | Group | Intcp Ctrl | Slp Ctrl | Median dist. Ctrl-Deming | n Ctrl | Median dist. Grp-Deming | n group | Est. diff. | CI 2.5% | CI 97.5% | Statistics Wilcoxon | p-value | Sig. | Eff. Size |
|---|---|---|---|---|---|---|---|---|---|---|---|---|---|---|
| Learning | CN-CL CNO during task | 110 | -0.704 | -3.60 | 168 | -23.2 | 40 | -21.6 | -31.7 | -12.3 | 1857 | 0.0000112 | *** | 0.305 |
| Consolidation | CN-CL CNO during task | -14.3 | 1.19 | 1.47 | 168 | -2.59 | 40 | 0.0355 | -9.81 | 9.99 | 3362 | 0.997 | n.s. | 0.000405 |
| Learning | CN-CL CNO after task | 110 | -0.704 | -3.60 | 168 | -8.14 | 32 | -5.61 | -18.4 | 7.99 | 2425 | 0.382 | n.s. | 0.062 |
| Consolidation | CN-CL CNO after task | -14.3 | 1.19 | 1.47 | 168 | 3.255 | 32 | 1.77 | -11.1 | 14 | 2755 | 0.825 | n.s. | 0.0158 |
| Learning | CN-VAL CNO during task | 110 | -0.704 | -3.60 | 168 | -4.28 | 40 | -0.967 | -10.9 | 9.26 | 3292 | 0.844 | n.s. | 0.0138 |
| Consolidation | CN-VAL CNO during task | -14.3 | 1.19 | 1.47 | 168 | -11.2 | 40 | -14.9 | -23.7 | -6.32 | 2225 | 0.000913 | *** | 0.23 |
| Learning | CN-VAL CNO after task | 110 | -0.704 | -3.60 | 168 | -1.53 | 32 | 1.27 | -12.2 | 14.4 | 2749 | 0.84 | n.s. | 0.0144 |
| Consolidation | CN-VAL CNO after task | -14.3 | 1.19 | 1.47 | 168 | -11.2 | 32 | -17.7 | -29.4 | -7.16 | 1742 | 0.00163 | ** | 0.223 |

Supplementary Table 10: Statistics for Fig 3g. Deming regression.

| Experiment | Value | ANOVA Group | p-value | Sig. |
|---|---|---|---|---|
| Footprint | Linear movement | F(3,31)=0.887 | 0.459 | n.s. |
| | Sigma | F(3,31)=0.103 | 0.958 | n.s. |
| | Alternate coefficient | F(3,31)=0.476 | 0.701 | n.s. |
| Horizontal Bar | Latency | F(3,33)=0.826 | 0.489 | n.s. |
| Vertical Pole | Latency | F(3,33)=0.968 | 0.42 | n.s. |
| Grid test | Latency | F(3,33)=0.305 | 0.822 | n.s. |

Supplementary Table 11: Statistics for SupFig 4abcd.

| Group | Day | Factor | ANOVA | p-value | Sig. | Group 1 | Group 2 | Statistic | CI 2.5% | CI 97.5% | p-value | Cohen's D | Sig. |
|---|---|---|---|---|---|---|---|---|---|---|---|---|---|
| CN-CL | 1 | Treatment | F(1,44)=1.443 | 0.236 | n.s. | | | | | | | | |
| | | Treatment:Moment | F(1,44)=2.457 | 0.124 | n.s. | | | posthoc | | | | | |
| | 4 | Treatment | F(1,44)=15.56 | <0.001 | *** | | | t-test | | | | | |
| | | Treatment:Moment | F(1,44)=16.594 | <0.001 | *** | OF1 (before SAL) | OF1 (before CNO) | 2.729 | 0.416 | 3.08 | 0.013 | 1.1 | * |
| | | | | | | OF2 (after SAL) | OF2 (after CNO) | -0.053 | -1.14 | 1.08 | 0.958 | -0.023 | n.s. |
| | 7 | Treatment | F(1,44)=2.483 | 0.122 | n.s. | | | | | | | | |
| | | Treatment:Moment | F(1,44)=2.31 | 0.136 | n.s. | | | | | | | | |
| CN-VAL | 1 | Treatment | F(1,36)=16.74 | <0.001 | *** | | | | | | | | |
| | | Treatment:Moment | F(1,36)=4.745 | 0.036 | * | OF1 (before SAL) | OF1 (before CNO) | -2.607 | -3.61 | -0.363 | 0.02 | -1.17 | * |
| | | | | | | OF2 (after SAL) | OF2 (after CNO) | -0.976 | -1.95 | 0.74 | 0.348 | -0.436 | n.s. |
| | 4 | Treatment | F(1,36)=0.197 | 0.66 | n.s. | | | | | | | | |
| | | Treatment:Moment | F(1,36)=10.455 | 0.003 | ** | OF1 (before SAL) | OF1 (before CNO) | -1.785 | -1.31 | 0.112 | 0.093 | -0.798 | n.s. |
| | | | | | | OF2 (after SAL) | OF2 (after CNO) | 0.889 | -0.62 | 1.53 | 0.386 | 0.397 | n.s. |
| | 7 | Treatment | F(1,36)=20.015 | <0.001 | *** | | | | | | | | |
| | | Treatment:Moment | F(1,36)=0.61 | 0.44 | n.s. | OF1 (before SAL) | OF1 (before CNO) | -1.794 | -1.79 | 0.142 | 0.09 | -0.802 | n.s. |
| | | | | | | OF2 (after SAL) | OF2 (after CNO) | -1.584 | -1.36 | 0.2 | 0.134 | -0.708 | n.s. |

Supplementary Table 12: Statistics for Sup Fig 4e.

| Group | Factor | ANOVA | p-value | Sig. | Group 1 | Group 2 | Statistic | CI 2.5% | CI 97.5% | p-value | Cohen's D | Sig. |
|---|---|---|---|---|---|---|---|---|---|---|---|---|
| CN-CL | Treatment | F(1,106)=0.416 | 0.52 | n.s. | | | | | | | | |
| | Speed | F(1,106)=116.649 | <0.001 | *** | | | | | | | | |
| | Treatment:Speed | F(1,106)=0.007 | 0.934 | n.s. | | | | | | | | |
| CN-VAL | Treatment | F(1,96)=17.495 | <0.001 | *** | | | | | | | | |
| | Speed | F(1,96)=132.095 | <0.001 | *** | | | | | | | | |
| | Treatment:Speed | F(1,96)=1.024 | 0.314 | n.s. | 5 rpm-SAL | 5 rpm-CNO | 0.876 | -23.3 | 53.3 | 0.402 | 0.392 | n.s. |
| | | | | | 10 rpm-SAL | 10 rpm-CNO | 1.21 | -17.1 | 56.5 | 0.257 | 0.541 | n.s. |
| | | | | | 15 rpm-SAL | 15 rpm-CNO | 3.81 | 22.6 | 82.8 | 0.002 | 1.7 | ** |
| | | | | | 20 pm-SAL | 20 rpm-CNO | 2.37 | 5.59 | 94.8 | 0.03 | 1.06 | * |
| | | | | | 25 rpm-SAL | 25 rpm-CNO | 2.04 | -2.58 | 58.8 | 0.069 | 0.914 | n.s. |

Supplementary Table 13: Statistics for Sup Fig 4f.

| Group | Time of treatment | Day | ANOVA Treatment | p-value | Sig. | ANOVA Treatment:Trial | p-value | Sig. |
|---|---|---|---|---|---|---|---|---|
| CN-CL | During task | 7 | F(1,18)=8.386 | 0.00963 | ** | F(6,108)=0.5056 | 0.803 | n.s. |
| | | 8 | F(1,18)=6.786 | 0.017 | * | F(6,108)=0.3734 | 0.894 | n.s. |
| | | 9 | F(1,18)=3.659 | 0.071 | n.s. | F(6,108)=0.5489 | 0.769 | n.s. |
| CN-VAL | During task | 7 | F(1,19)=9.661 | 0.00579 | ** | F(6,114)=1.223 | 0.3 | n.s. |
| | | 8 | F(1,19)=3.106 | 0.094 | n.s. | F(6,114)=0.5508 | 0.768 | n.s. |
| | | 9 | F(1,19)=4.405 | 0.049 | * | F(6,114)=1.807 | 0.103 | n.s. |

Supplementary Table 14: Statistics for Fig 4ac.

| Time of treatment | Group | Measure | n | Comparison | Estimate | Test | Alternative | Statistic | p-value | Sig. | 95% CI high | 95% CI | Effect size |
|---|---|---|---|---|---|---|---|---|---|---|---|---|---|
| During task | CN-CL SAL-CNO | daily start | 10 | day 7 vs <day8 day9> | -76.2 | Wilcoxon | two.sided | 8 | 0.0488 | * | -119 | -23.5 | 0.629 |
| | | daily end | 10 | | -72.2 | | | 5 | 0.0195 | * | -124 | -14.1 | 0.725 |
| | CN-CL CNO-SAL | daily start | 10 | | 49 | | | 55 | 0.00195 | ** | 18.9 | 73.9 | 0.886 |
| | | daily end | 10 | | 39.1 | | | 45 | 0.084 | n.s. | -1.59 | 83.5 | 0.564 |
| | CN-VAL SAL-CNO | daily start | 11 | | -65.6 | | | 0 | 0.000977 | *** | -106 | -26.2 | 0.885 |
| | | daily end | 11 | | -73.3 | | | 10 | 0.042 | * | -128 | -1.48 | 0.617 |
| | CN-VAL CNO-SAL | daily start | 10 | | 19.6 | | | 39 | 0.275 | n.s. | -12.9 | 40.4 | 0.371 |
| | | daily end | 10 | | 14.9 | | | 39 | 0.275 | n.s. | -15 | 49 | 0.371 |

Supplementary Table 15: Statistics for Fig4bd.

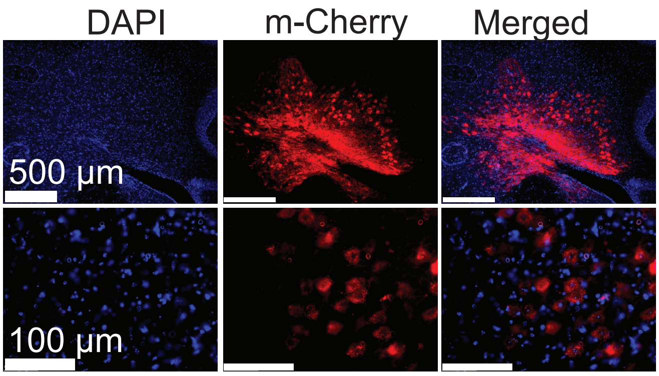


**Supplementary Figure 1.** DAPI positive neurons expressing hM4Di-DREADD-mCherry in cerebellar nuclei.

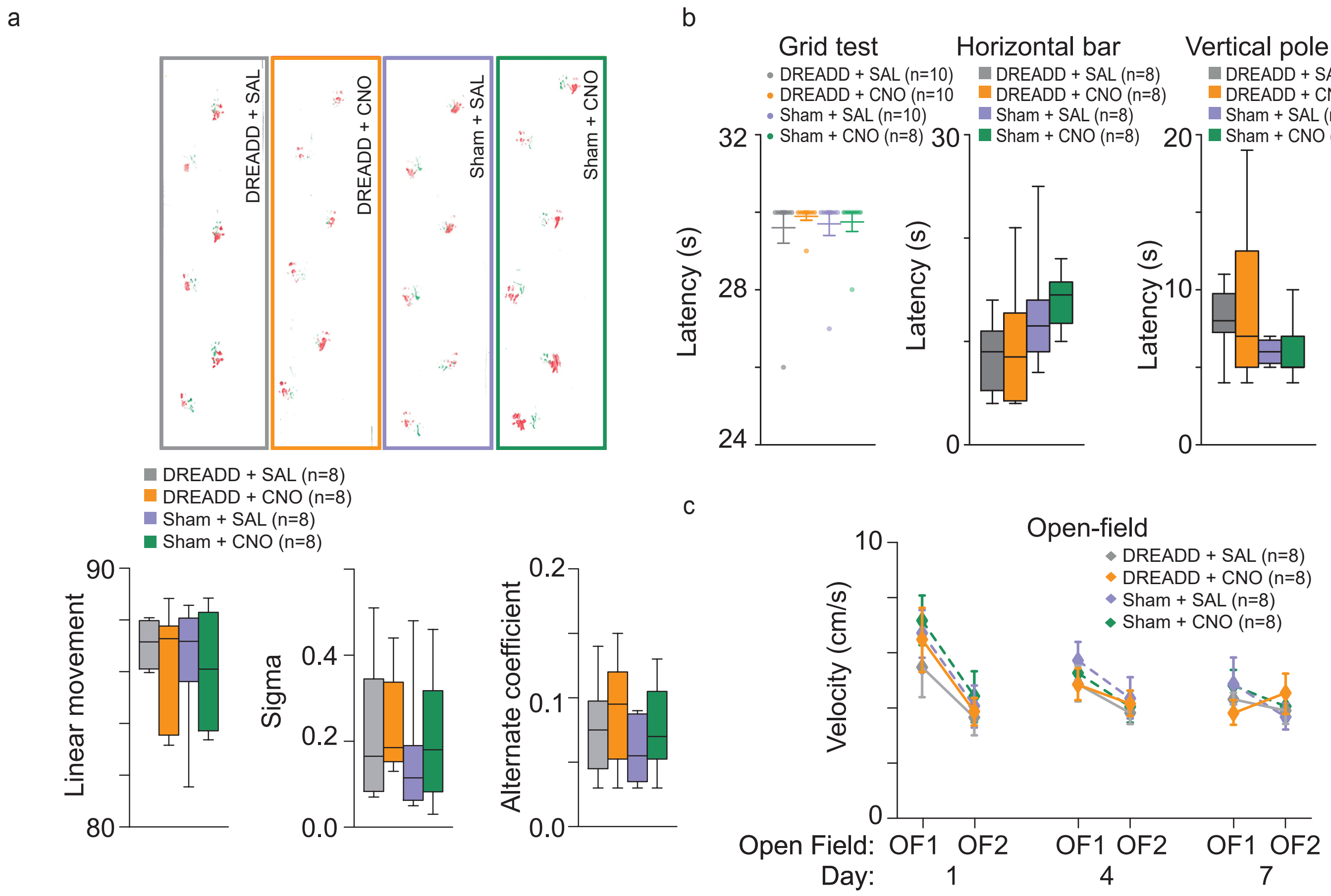


**Supplementary Figure 2. Cerebellar nuclei inhibition did not affect execution and fatigue, locomotion, motor coordination, balance and strength. a**) Footprint patterns were quantitatively assessed for 3 parameters as shown on representative footprint patterns for all experimental groups. Three parameters are represented graphically: linear movement (bottom left), sigma (bottom middle) and alternation coefficient (bottom right). **b**) tests of strength and coordination. Grid test: latency reflecting the time before falling from the grid. 30 seconds of cut-off of was established as the maximum latency (dotted line on figure). Horizontal bar: latency to cross the horizontal bar (balance beam test) for all experimental groups. Vertical pole: latency to reach home cage in vertical pole test for all experimental groups. **c**) Locomotor activity (Velocity) in DREADD and non-DREADD (Sham) injected mice after CNO or SAL injection during open-field sessions before (OF1) and after (OF2) rotarod for experimental days 1, 4 and 7. *n* indicates the number of mice. Boxes represent quartiles and whiskers correspond to range; points are singled as outliers if they deviate more than 1.5 x interquartile range from the nearest quartile. No posthoc significant difference was found for the data presented in this figure (see Supplementary Tables 6-7 for detailed statistics).

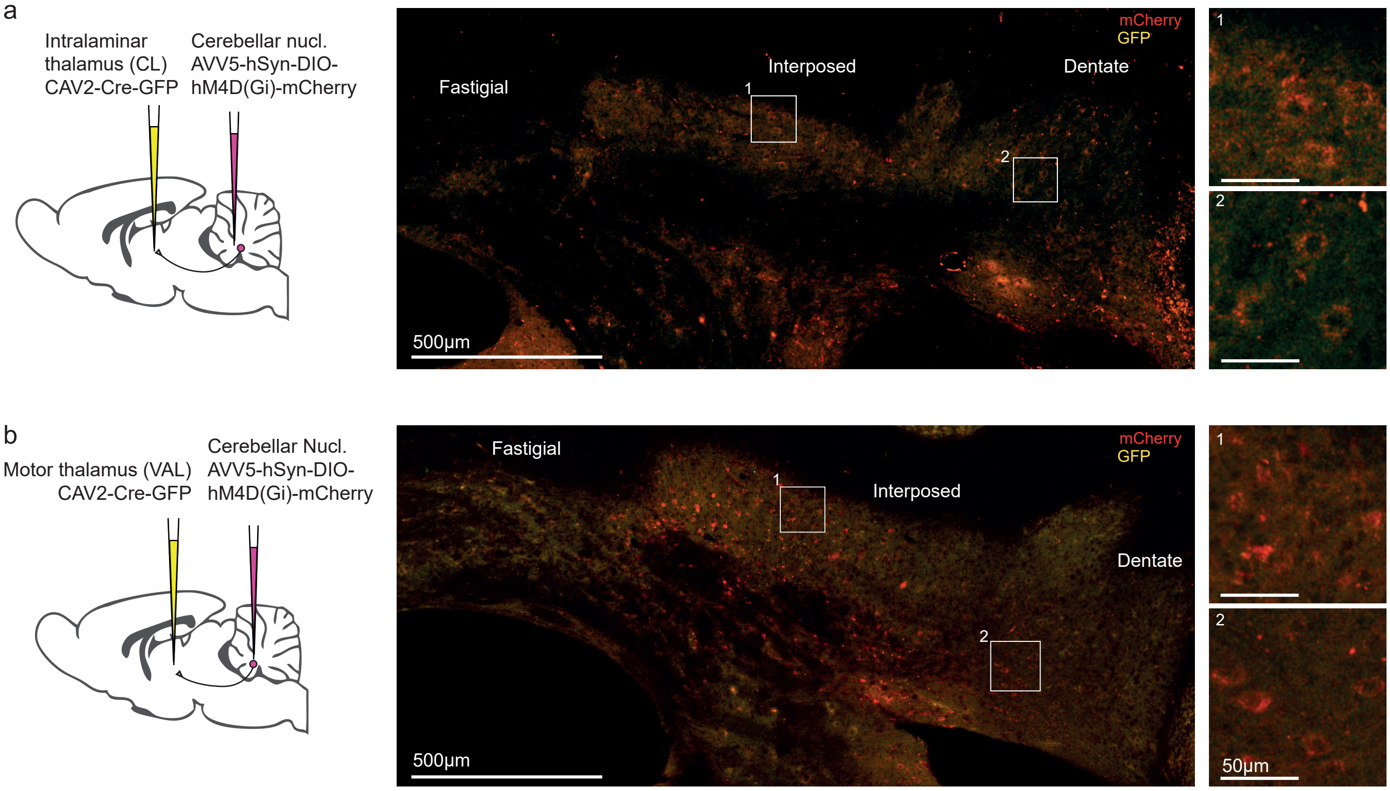


Supplementary Figure 3. Expression of hM4D(Gi)-mCherry in the cerebellar nuclei following CL and VAL injections. a) left: schematics of the experiment for CN-CL groups, right: distribution of labeled neurons. b) same as a for CN-VAL groups.

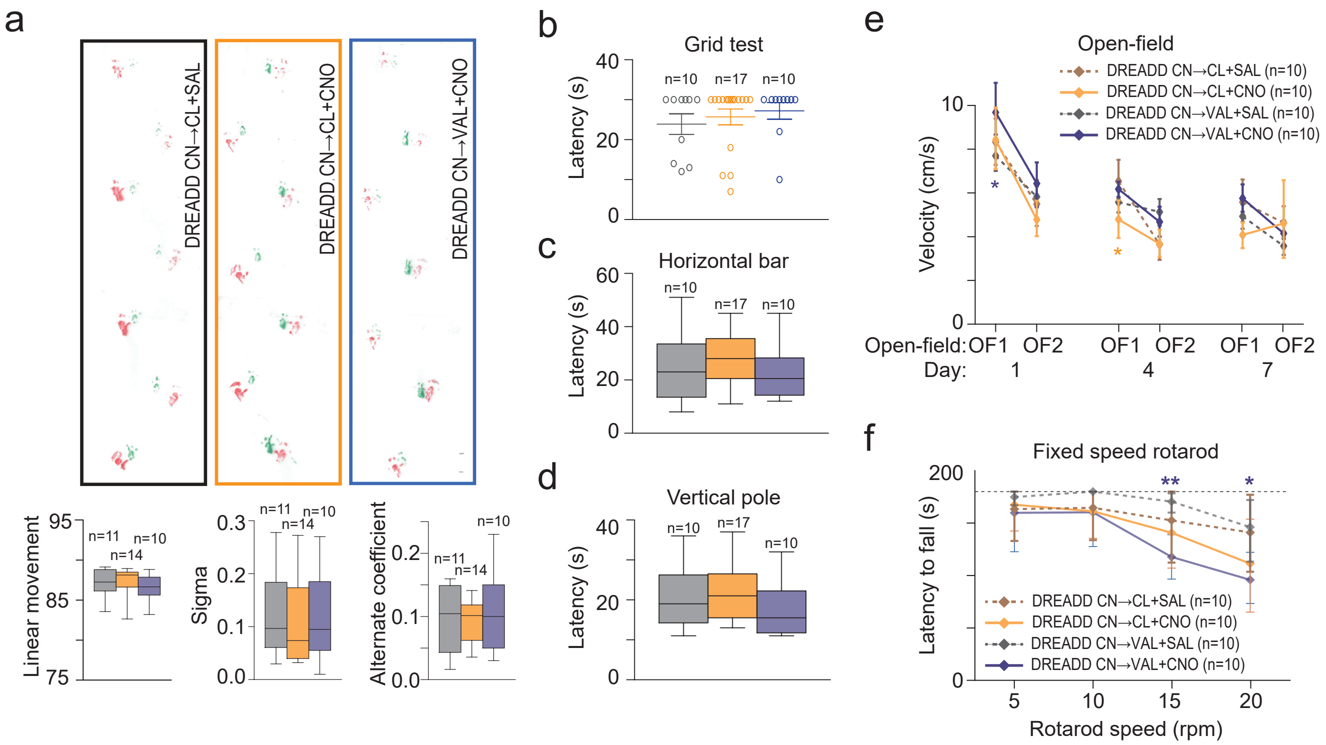


**Supplementary Figure 4. Inhibition of CN-CL or CN-VAL by 1mg/kg CNO does not affect execution and fatigue, locomotion, motor coordination, balance and strength**. **a**) Footprint patterns were quantitatively assessed for 3 parameters as shown on representative footprint patterns (top) for all experimental groups. Three parameters are represented graphically: linear movement (bottom left), sigma (bottom middle and alternation coefficient (bottom right). **b**) Latency reflecting the time before falling from the grid. 30 seconds of cut-off of was established as the maximum latency (dotted line on figure). **c**) Latency to cross the horizontal bar (balance beam test) for all experimental groups. **d**) Latency to reach home cage in vertical pole test for all experimental groups. **e**) Locomotor activity (Velocity) in DREADD injected mice after CNO or SAL injection during open-fields sessions before (OF1) and after (OF2) rotarod for day 1, 4 and 7 (**$p<0.01$ paired t-test OF1 vs OF2). **f**) Latency to fall during fixed speed rotarod (5, 10, 15, 20 r.p.m.) for all experimental groups. One-way repeated measure ANOVA was performed on averaged values for each speed steps in each experimental group followed by a Tukey Posthoc pairwise comparison (*$p<0.05$, **$p<0.01$ for CN->VAL group CNO vs SAL). CN, cerebellar nuclei, CL, centrolateral thalamus; VAL, ventral anterior lateral thalamus. *n* indicates the number of mice. Boxes represent quartiles and whiskers correspond to range; points are singled as outliers if they deviate more than 1.5 x interquartile range from the nearest quartile. (See Supplementary Tables 11-13 for detailed statistics)

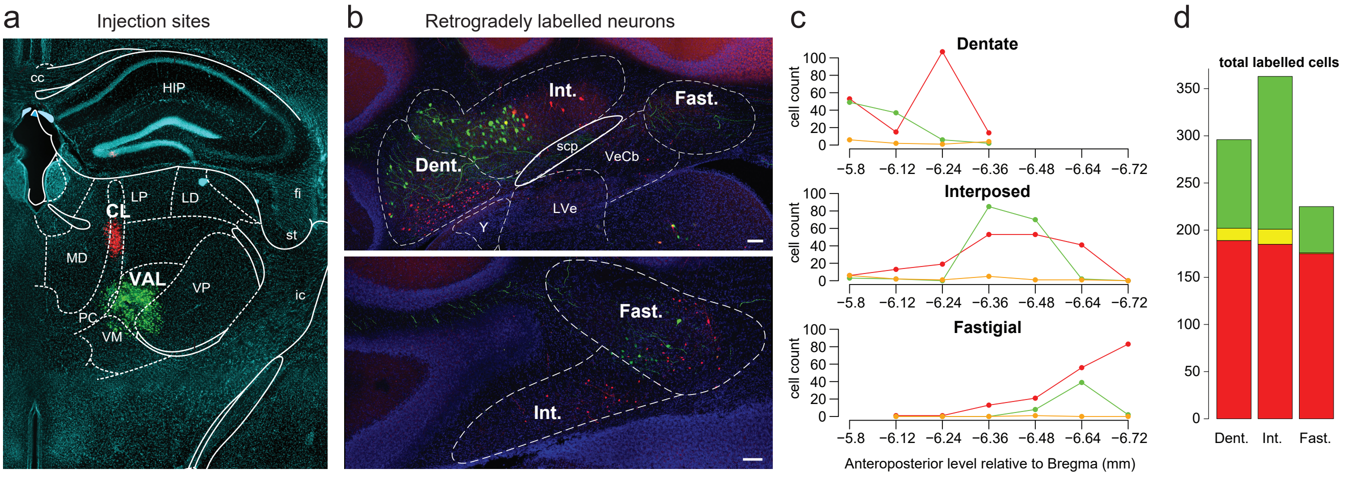


**Supplementary Figure 5: retrograde infections from the CL and VAL differentially label cerebellar nuclei populations.** a. injections sites targeted at the VAL (AAVrg-CAG-GFP) and CL (AAVrg-CAG-mCherry). b. example of retrogradely infected neurons at two different levels of the cerebellar nuclei. c. summary of neuronal counts at 7 different anteroposterior levels (cumulated from 2 mice, where counts at each level and each cerebellar nuclei exhibit high correlation r=0.92). Red infection from the CL, green infection from VAL, orange: infection from both structures. d. summary of total cell counts per nucleus.

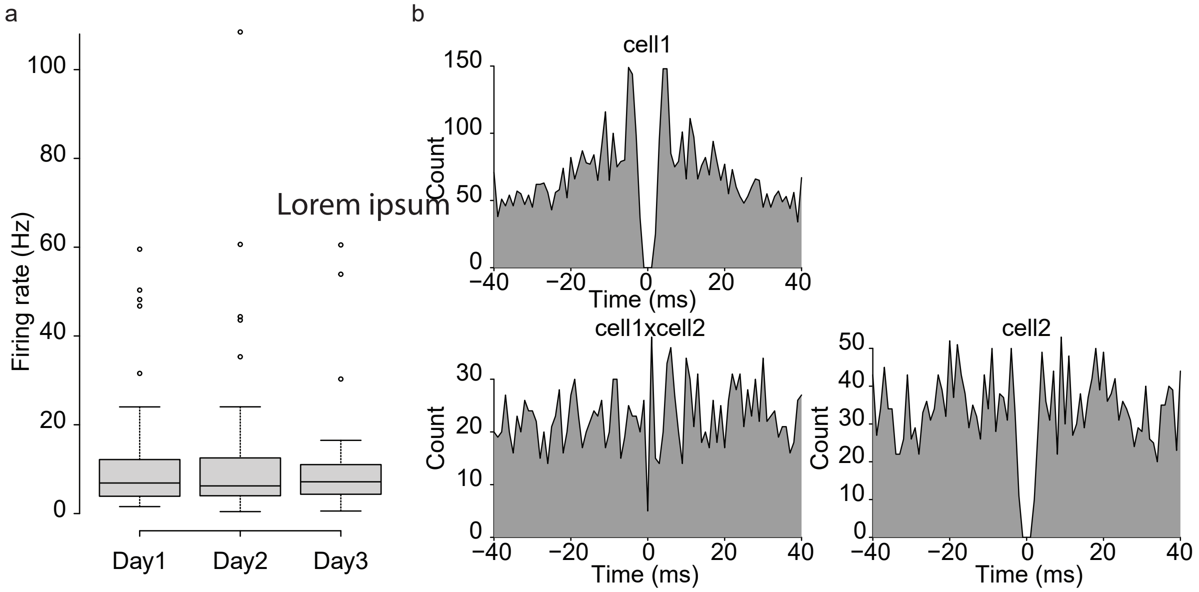


Supplementary Figure 6: extracellular recordings in the CN. a. similarity of initial openfield firing rates for cerebellar units isolated on successive days. b. example of auto- and cross-correlogram from stimultaneously recorded units.
